# The vaccinia virus chondroitin sulfate binding protein drives host membrane curvature to facilitate fusion

**DOI:** 10.1101/2022.03.10.483741

**Authors:** Laura Pokorny, Jemima J. Burden, David Albrecht, Rebecca Bamford, Kendra E. Leigh, Pooja Sridhar, Timothy J. Knowles, Yorgo Modis, Jason Mercer

## Abstract

Virus binding serves to define virus tropism and species specificity^1^. Virus binding proteins are classically considered as facilitators of cell surface attachment prior to receptor engagement and virus internalization. For efficient entry vaccinia virus (VACV) – the prototypic poxvirus - relies on four binding proteins and an eleven-protein entry fusion complex (EFC)^2^. We recently demonstrated that VACV binding and fusion proteins are organized into distinct functional domains, with localization of EFC proteins to virion tips directly influencing membrane fusion activity^3^. However, the relationship between virus binding protein distribution, virion binding orientation and subsequent membrane fusion remain unexplored. Here, we show that virus binding proteins guide side-on virion binding and promote curvature of the host membrane towards EFC-containing virion tips to facilitate virus fusion. Using a cell-derived membrane-bleb model system together with super-resolution and electron microscopy we found that side-bound VACV virions induce membrane invagination in the presence of low pH. Repression or deletion of individual binding proteins revealed that three of four contribute to binding orientation, amongst which the chondroitin sulphate binding protein, D8, is required for host membrane bending. Consistent with low-pH dependent macropinocytic entry of vaccinia virus^4,5^, loss of D8 prevents virion-associated macropinosome membrane bending, disrupts fusion pore formation and infection kinetics. Our results extend the role of viral binding proteins from mere attachment factors to active participants in successful viral membrane fusion and further illustrate the influence of virus protein architecture on successful infection.

---

Infectious mature virions (MVs) of all poxviruses, including variola (the causative agent of smallpox), monkeypox and the smallpox vaccine, vaccinia virus (VACV), encode 4 distinct binding proteins and an 11-protein entry fusion complex (EFC)^6^. Each of the 11 EFC proteins is needed for membrane fusion competence^2,7^, while the 4 reported binding proteins A26, A27, D8 and H3 differentially contribute to MV attachment^8–12^. The A26 protein has been shown to bind extracellular laminin, serve as a fusion repressor and its exclusion from virions, to impact virion binding^12–14^. A27, H3 and D8 are glycosaminoglycan (GAG) binding proteins reported to interact with heparin sulfate (A27, H3) or chondroitin sulfate (D8), respectively^9–11^. While soluble A27, H3 and D8 each interfere with virion binding, only deletion of D8 was shown to significantly impact GAG-mediated cell surface attachment of MVs^9–11^. Collectively, various reports indicate that VACV GAG usage varies with cell type, virus strain and experimental condition^15–18^. Due to this complexity, the intricacies and individual contribution of these proteins to VACV MV cell surface binding remain poorly understood.

We recently showed that the VACV membrane is organised into distinct functional domains; with EFCs polarised to the tips and binding proteins A27 and D8 relegated to the sides^3^. A27-dependent EFC polarisation was found to be critical for tip-oriented fusion, and localization of D8 to correlate with a side-on virion binding bias^3,19^. This begged the questions: Which of the four viral binding proteins contribute to VACV binding orientation and how does this correlate or contribute to tip-oriented fusion?

Traditional membrane model systems such as liposomes and giant unilamellar vesicles (GUVs) have been invaluable for investigating binding, uptake and fusion of several viruses^19–22^. However, as VACV binding relies on the presence of multiple cell surface proteoglycans, investigation in these lipidic systems is inefficient and perhaps biologically irrelevant^23^. To this end we established a minimal model system based on cell-derived membrane blebs^24^ to facilitate the quantification of large numbers of binding events in both fluorescence- and electron microscopy (EM)- based imaging experiments (Supplementary Fig. 1). Using a fluorescent-recombinant VACV (EGFP-A4^5^) we found that blebs could be used for analysis of virus binding (Fig. 1a), and when coupled with our established octadecylrhodamine (R18)-dequenching assay^3,23^, pH-dependent hemi-fusion (Fig. 1b). Use of the fusion neutralizing anti-L1 antibody (7D11) confirmed that the observed low pH-mediated dequenching was due to virus-bleb fusion and not non-fusogenic dye transfer^23,25^ (Fig. 1b).

**Figure 1.**
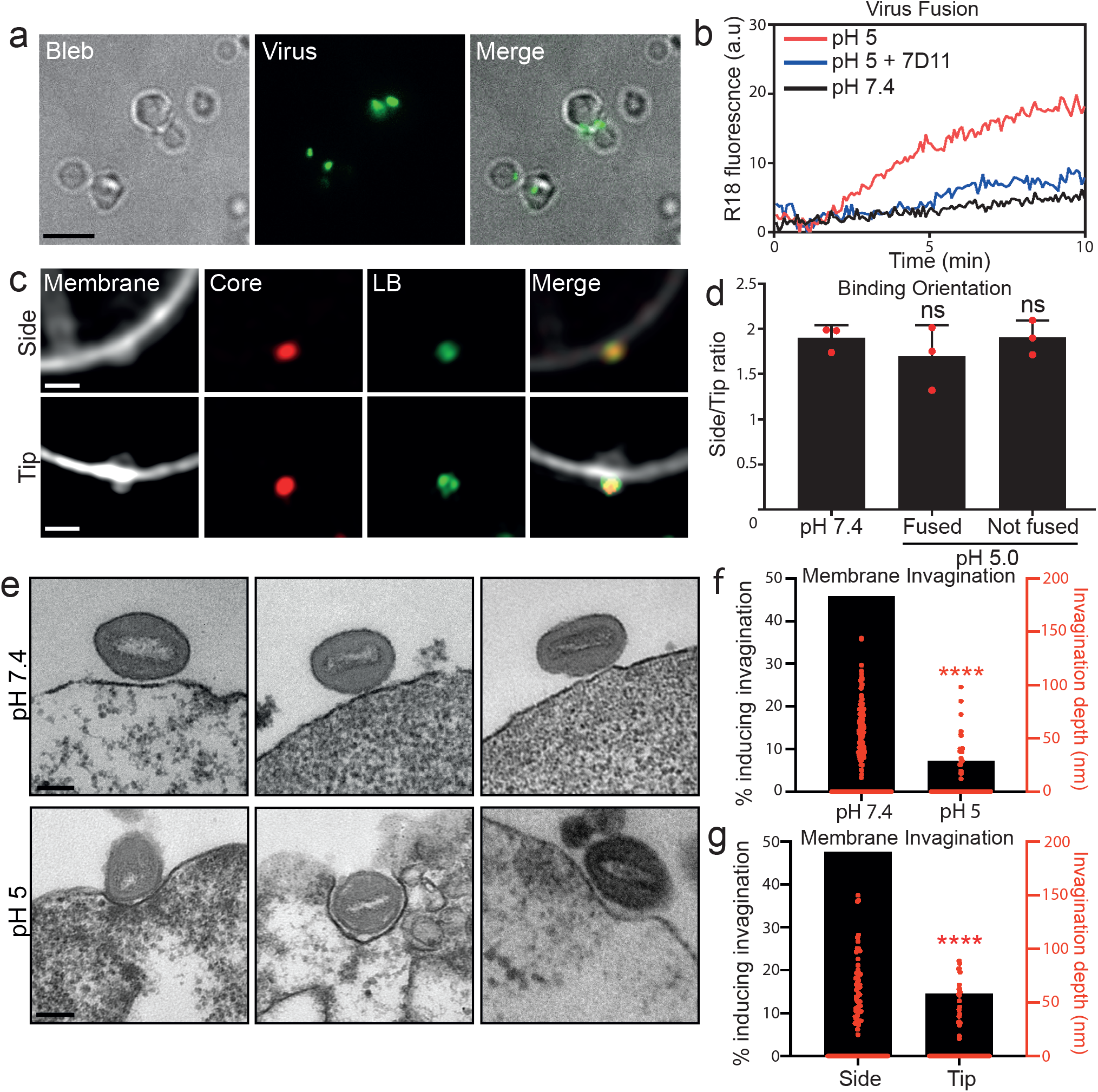
VACV binds to, fuses with and mediates curvature of membrane blebs. **a.** Blebs incubated with A4-EGFP virus at 4 °C for 1 hr, washed and imaged. Scale bar, 5um. **b.** VACV hemifusion rates (R18 dequenching assay) with blebs at pH 7.4 or pH 5.0. 7D11 fusion neutralizing antibody was used as a positive control. Fluorescence was normalised to the initial value and fully de-quenched value upon TX-100 addition. **c.** Examples SIM images of mCherry-A4 EGFP-F17 VACV bound to blebs in a side-on (upper row) and tip-on (lower row) orientation. Scale bar = 1 μm. **d.** Quantification of side/tip binding ratio at different pH’s and fusion states **e.** TEM images of VACV bound to HeLa cell derived blebs at pH 7.4 (top row) or pH 5.0 (bottom row). Scale bar = 100 nm. **f.** Quantification of membrane invagination depth under virions at pH 7.4 and pH 5.0. **g.** Quantification of side-on vs. tip-on bound virion invagination. For **d** (n=150), **f** (n=50) and **g** (n=100) data are mean ± standard deviation (SD) of biological triplicates. Statistical analysis was performed using unpaired two-tailed *t*-tests (\*\*\*\**P* < 0.0001; ns, not significant (*P* > 0.05)).

Having shown that our bleb model system supports virus binding and fusion, we turned our attention to accurately identifying and quantifying VACV binding orientation during fusion. Using scanning- and transmission-EM (TEM) respectively we recently reported that VACV virions preferentially bind on their sides and undergo low-pH dependent fusion at their tips^3^. This suggested to us that membrane-bound virus may undergo reorientation in response to low pH bringing the virion tips into contact with the cell membrane. To test this, recombinant virions harbouring a mCherry-tagged core protein and an EGFP-tagged lateral body protein (mCherry-A4 F17-EGFP)^26,27^ were loaded with R18 dye, bound to blebs and subjected to neutral (7.4) or low (5.0) pH treatment. Blebs bound with a single virion were visualized by Structured Illumination Microscopy (SIM) (Fig. 1c). Using core elongation and lateral body separation as parameters, virion binding orientation was plotted as side/tip ratio (Fig 1c,d). The pH 5.0 samples were also scored for virion hemi-fusion as evidenced by R18-dequeching into the bleb membrane (Fig. 1c,d). In all cases virions showed preferential side binding with no significant changes in binding orientation observed upon low pH treatment or subsequent hemi-fusion (Fig.1d). Intriguingly, these results suggested that despite the localization of the EFC to virion tips, VACV does not need to be bound on its end to undergo fusion with host membranes.

To reconcile this, we used TEM to further investigate the binding interaction between virions and blebs at pH 7.4 and pH 5.0. As expected, under both conditions, virions bound preferentially on their sides. While we observed no discernible changes in the underlying bleb membrane at pH 7.4 (Fig. 1e; top row), at pH 5.0 invagination of the bleb membrane occurred specifically under bound virions (Fig. 1e, bottom row). Quantification indicated that at pH 7.4, only 7.1% of virions were found in invaginations between 11 and 72.2 nm deep (average depth = 36 nm), while at pH 5.0, 46% of virions resided in invaginations ranging from 13-144 nm deep (average depth =60 nm) (Fig. 1f). When the pH 5.0 samples were scored by virion binding orientation, 48% of side bound virions were found in invaginations as opposed to 15% of tip bound virions (Fig. 1g). Collectively these results confirm that VACV binds to host membranes in a side-on orientation, indicate that binding orientation is not altered by low pH or the induction of hemi-fusion, and suggest that when bound on their side VACV virions manipulate the curvature of cellular membranes in response to low pH.

That the VACV membrane is organized into functional binding and fusion domains^3^ we reasoned that VACV binding protein(s) may be responsible for the induction of host membrane curvature. Having shown that VACV binding proteins A27 and D8 reside at the sides of virions^3^, we first investigated the localization of the two remaining binding proteins, A26 and H3^11,12^. For this, we generated recombinant EGFP core (EGFP-A4) viruses that express HA-tagged versions of A26 or H3. Using SIM and the single-particle averaging software VirusMapper^27^, we mapped the localization of A26 and H3 on VACV virions. Virions immuno-stained for A27 and D8 were included for comparison. VirusMapper models and determination of the polarity factor indicated that like A27 and D8, A26 and H3 are largely relegated to the sides of virions (Fig. 2a, Supplementary Fig. 2a). We confirmed these results using Stochastic Optical Reconstruction Microscopy (STORM), a higher resolution technique that also suggested that A26 and H3 reside in the VACV membrane as clusters akin the A27 and D8 (Supplementary Fig. 2b;^3^).

**Figure 2.**
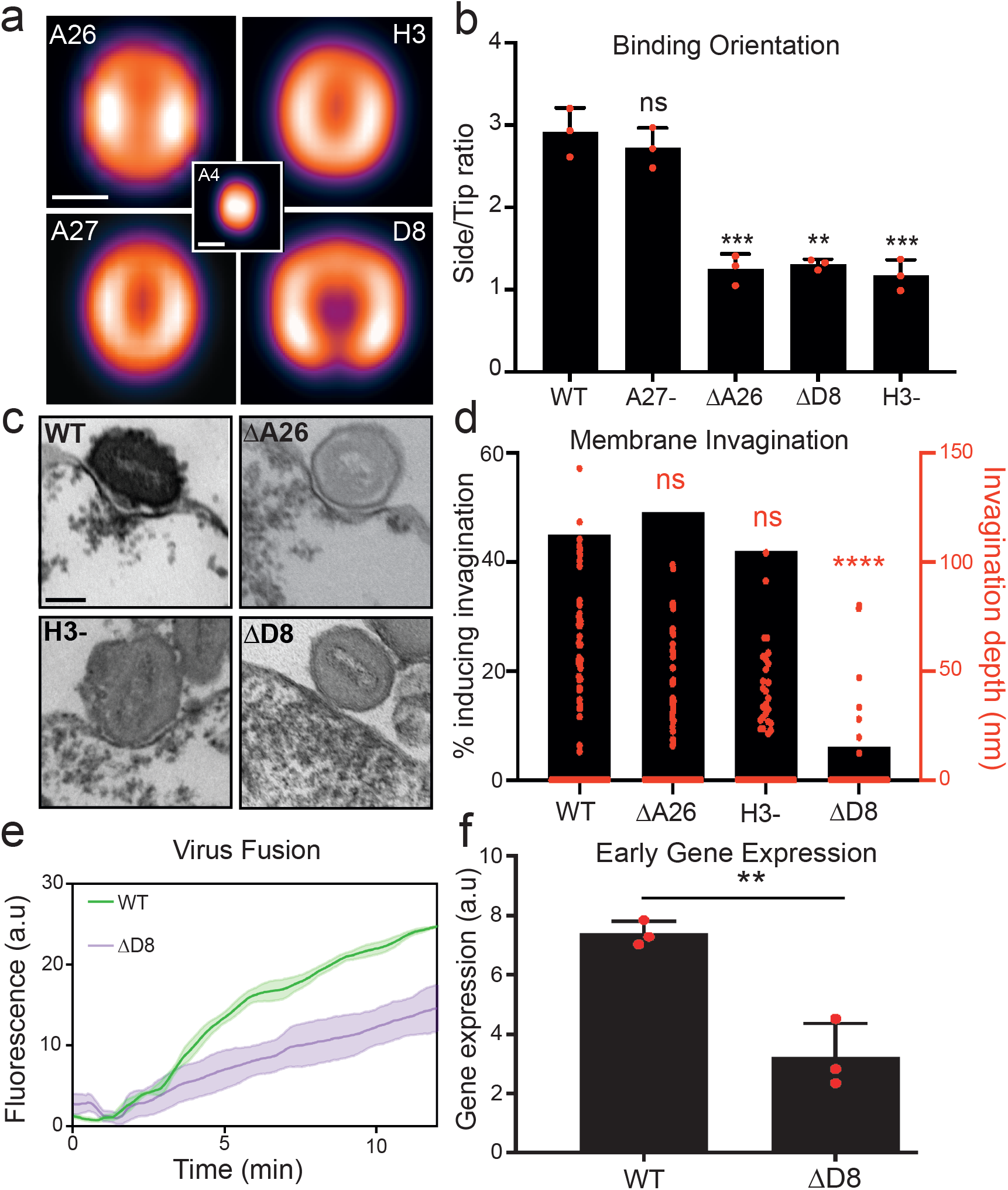
VACV binding protein D8 is required for host membrane invagination and fusion. **a. VirusMapper** localisation models of VACV binding proteins. EGFP-A4 core (center) was used to correlate virion orientation. Models are representative of n>180 virions. Scale bar, 200 nm. **b.** Analysis of side/tip binding ratio of WT and binding protein mutants on blebs. **c.** TEM images of WT and binding protein mutants bound to HeLa blebs at pH 5.0. Scale bar, 100 nm. **d.** Quantification of percent invagination and invagination depth of binding protein mutants at pH 5.0. **e.** Hemifusion rates of WT and ΔD8 virions on HeLa cells using the R18 dequenching assay. **f.** Comparison of WT and ΔD8 early gene expression at 2 hpi by RT-qPCR of early gene C11R. For **b** (n=100) and **d** (n=100), **e** and **f** data are means ± SD of biological triplicates. Statistical analysis was performed using unpaired two-tailed *t*-tests (\*\*\*\**P* < 0.0001; \*\*\**P* < 0.001; \*\**P* < 0.01; \**P* < 0.05; ns, not significant (*P* > 0.05)).

To gain a comprehensive assessment of the relative contribution of these four proteins to VACV binding and binding orientation we generated EGFP-core versions of inducible A27 and H3 VACV recombinants, as well as VACV strains in which A26 or D8 have been deleted. We compared the binding capacity of WT virions to those lacking A26 (ΔA26), A27 (A27-), D8 (ΔD8) or H3 (H3-). For this, equal numbers of virions were adsorbed to cells and binding quantified by flow cytometry for EGFP signal. Loss of A27 showed no impact on VACV binding, while loss of A26, D8 or H3 reduced VACV binding by 66, 47, and 24%, respectively (Supplementary Fig. 2C). Results from our bleb binding orientation assay (Fig. 1c,d) showed similar results: A27-virions preferentially bind side-on like WT virions, while a significant proportion (2.2 fold) of ΔA26, DD8 and H3-virions adapt a tip-on binding orientation (Fig. 2b). These results suggest that despite its ability to bind heparin sulfate^8,10^, A27 plays no apparent role in VACV binding. They also demonstrate that A26, D8 and H3 each contribute to virion side-on binding orientation, and that A26 contributes most to cell surface attachment, followed by D8 and H3.

Given these results, it reasons that A26, D8, H3, or a combination thereof contribute to the observed low pH-dependent membrane invagination. Using TEM we compared the membrane invagination capacity of WT, ΔA26, ΔD8 and H3-virions on blebs at pH 5.0. While we could readily observe WT, DA26 and H3-virions residing in invaginations, that vast majority of DD8 virions were associated with unaltered bleb membranes (Fig. 2c) resembling WT virion binding at pH 7.4 (Fig. 1e; top row). Quantification showed that the percentage of WT, ΔA26 and H3-virions found within invaginations was similar, ranging between 42-49%, as opposed to only 6% of ΔD8 virions (Fig. 2d; black bars). This was consistent with a significant decrease in the average invagination depth of DD8 virions relative to WT, ΔA26 and H3-virions (Fig. 2d; red data points). Taken together, these results show that A26, H3 and D8 each contribute to VACV cell surface binding differentially. While all three proteins contribute to binding orientation to a similar degree, A26 appears to be most important for attachment, and D8 for a newly uncovered role in low pH dependent membrane invagination.

We next sought to determine the relevance of low pH dependent, D8-mediate membrane invagination. Given the distinct spatial distribution of the viral binding proteins to the sides and fusion proteins to the tips of VACV virons^3^, we hypothesized that this membrane “curving” activity of D8 could serve to bring the host membrane into contact with the virus fusion machinery under low pH conditions, such as those found in late macropinosomes from which VACV fuses^4,23,28^. If correct, we would expect VACV fusion activity to be impacted by the loss of D8. To investigate this, we compared WT and ΔD8 hemifusion rates on HeLa cells using our R18 dequenching assay.

ΔD8 hemifusion was found to be reduced by 1.7-fold relative to WT virus (Fig. 2e). This result correlated with a 2.3-fold reduction in early gene expression - the earliest read-out of core entry into the cytoplasm - as determined by real time quantitative PCR (RT-qPCR) for the canonical early gene, C11R (Fig. 2f). These results demonstrate that loss of D8, and its low pH-dependent membrane invagination activity, impacts VACV fusion efficiency and the kinetics of early gene expression.

VACV can enter cells by direct fusion at the plasma membrane^17,25^ or via low pH-dependent macropinocytosis^5,29,30^. Therefore, we wanted to determine if D8-mediated invagination occurs at the limiting membrane of macropinosomes during VACV entry. As VACV resident time in macropinosomes is relatively short^28^ making it difficult to capture in significant numbers, we took advantage of an A28-EFC mutant VACV which can undergo hemifusion, but not full fusion^7,31^. Importantly, using EM we first confirmed that A28-virions induce pH-dependent invagination of blebs akin to WT virions (WT:40% vs A28-:42%) (Fig. 3a,b).

**Figure 3.**
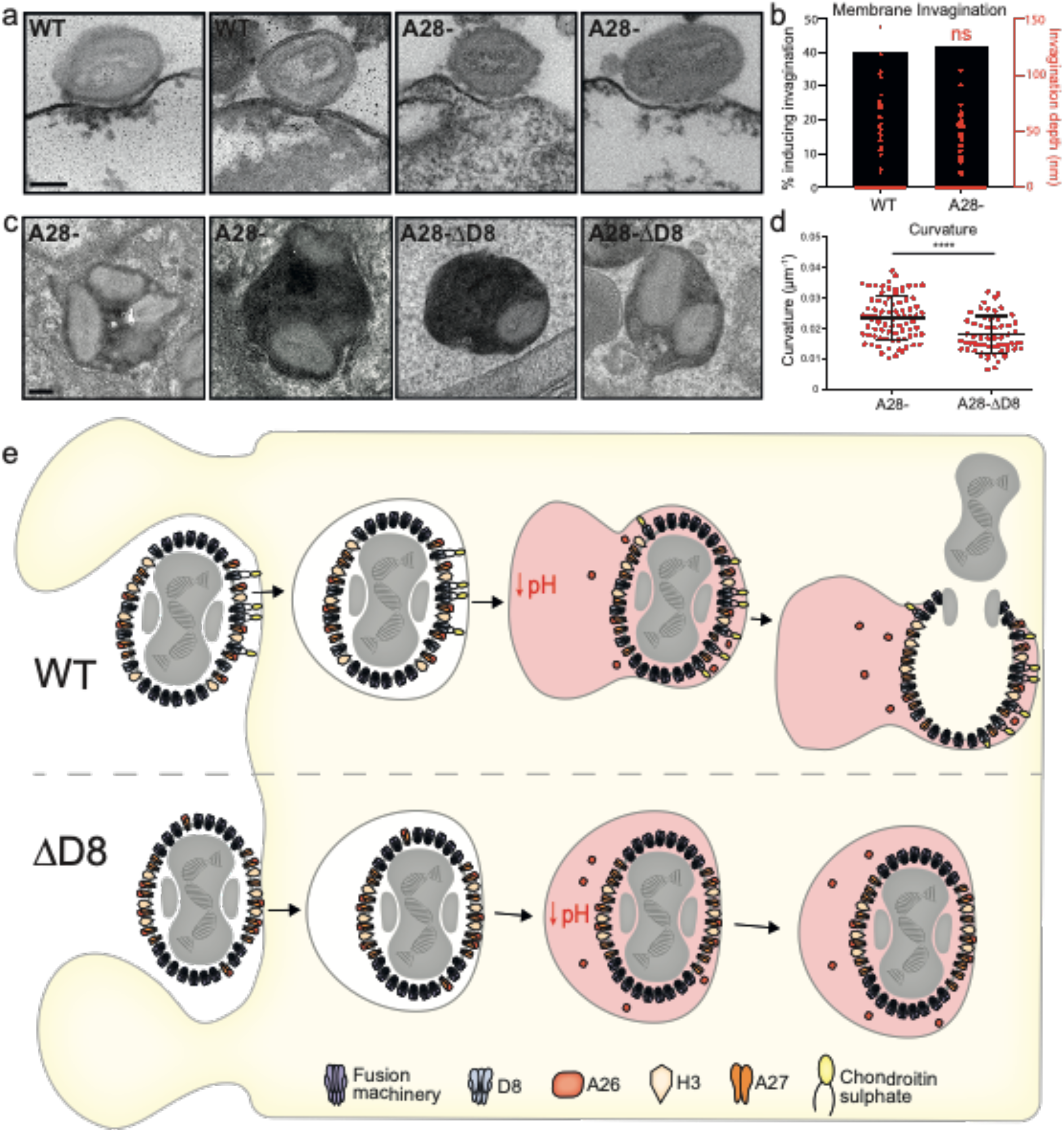
D8-mediated invagination is a critical component of VACV low pH-dependent fusion. **a.** Representative TEM images of WT and A28-virions bound to HeLa blebs at pH 5.0. Scale bar, 100 nm. **b.** Quantification of WT and A28-virion % invagination and invagination depth comparing under pH 5.0 conditions (from a). **c.** Representative TEM images of A28- or A28-ΔD8 virions within macropinosomes. Scale bar, 100 nm. **d.** Quantification of curvature of the macropinosome membrane around individual A28- or A28-ΔD8 virions (from f) using the Fiji plugin Kappa. **e.** Model of low-pH mediated D8-mediated macropinosome membrane curvature and VACV fusion. For **b** (n=100 virions/condition) and **d** (n=80 virions/condition) data are means ± SD of biological triplicates. Statistical analysis was performed using unpaired two-tailed *t*-tests (\*\*\*\**P* < 0.0001; \*\**P* < 0.01; ns, not significant (*P* > 0.05)).

We then generated an A28-ΔD8 virus, confirmed that it did not express D8 (Supplementary Fig. 3) and used this in combination with a TEM horseradish peroxidase (HRP) entry assay to investigate the association of WT A28-ΔD8 virions with the macropinosome membrane. The HRP was used to increase the electron density within endosomes, making them easily visible by TEM, and to ensure that only endocytic events were analysed^32,33^. The TEM images showed notable qualitative differences in both macropinosome shape and virion-macropinosome association between VACV A28- and A28-ΔD8 samples (Fig. 3c). A28-virions were tightly wrapped causing the macropinosome to take on the shape of the virions within. Of note, the limiting membrane of macropinosomes were often found in contact with A28-virion tips. In contrast A28-ΔD8 containing macropinosomes remained circular with little evidence of virion-mediated membrane deformation. To analyse the curvature of the macropinosome membrane induced by single virions, Kappa, a Fiji plugin which measures curvature using B-splines, was used^34^. Only virions in contact with the limiting macropinosome membrane, in a side-on orientation, were considered for analysis to avoid skewing the data with high curvature values around the tip of A28-virions. Using these parameters, a 1.3-fold drop in macropinosome membrane curvature around virions was seen in the absence of D8 (Fig. 3d).

These results are consistent with the model shown in Figure 3e, whereby WT VACV virions bind to the cell surface on a side on orientation via A26-lamanin, D8-chondrotin sulfate, and H3-heparan sulfate interactions before being macropinocytosed. As the macropinosome matures lowering the pH within, the fusion repressor A26 is removed, and D8-mediated macropinosome membrane curvature is triggered. This serves to bring the limiting membrane of the macropinosome in contact with the fusion machinery, located at virion tips, thereby increasing the likelihood of productive fusion. When virions lacking D8 (ΔD8) are macropinocytosis their inability to drive membrane invagination - despite side on binding and low-pH mediated removal of A26 - slows the rate virion-target membrane fusion. Taken together our results strongly suggest that a critical component of fusion from macropinosomes is VACV low pH dependent D8-mediated membrane invagination.

Next, we turned to the cellular factors involved in this phenomenon. Like many viruses, VACV uses cellular glycosaminoglycans, such as heparan sulfate (HS) and chondroitin sulfate (CS), as cell surface attachment factors^1,35^. As D8 is a well-defined CS-binding protein^9,36^, it seemed likely that the D8-CS interaction was the driving force behind D8 membrane invaginating activity. To test this, we took advantage of WT L929 cells that display both HS and CS (HS+CS+), and the L929 derivatives gro2c which display only CS (HS-CS+), and sog9 which display neither HS or CS (HS-CS-)^37,38^.

Confirming the importance of CS in VACV attachment, 39% less A4-EGFP VACV virions bound to cells lacking CS than to cells containing both HS and CS, or cells lacking HS alone (Supplementary Fig. 4a). To investigate the role of CS in D8-mediated invagination we generated HS+CS+ and HS-CS- blebs - confirmed that VACV could bind to them (Supplementary Fig. 4c) - and compared invagination under WT virions at pH 5.0. Membrane invagination under virions was evident in HS+CS+ blebs and absent in HS-CS- blebs (Fig. 4a). Quantification indicated that 39% of virions could induce invagination of HS+CS+ blebs, as opposed to just 8.5% in HS-CS- blebs, this was accompanied by a significant reduction in invagination depth (Fig. 4b) Consistent with the decreased rate of hemifusion observed with ΔD8 virus (Fig. 3b), the rate of VACV hemifusion on HS-CS- cells was reduced by 1.9-fold relative to HS+CS+ cells (Fig. 4C). Together with our binding experiments, these results indicate that CS - through its interaction with D8 - is important for both virus attachment to the cell surface and the rate of low pH-dependent VACV fusion.

**Figure 4.**
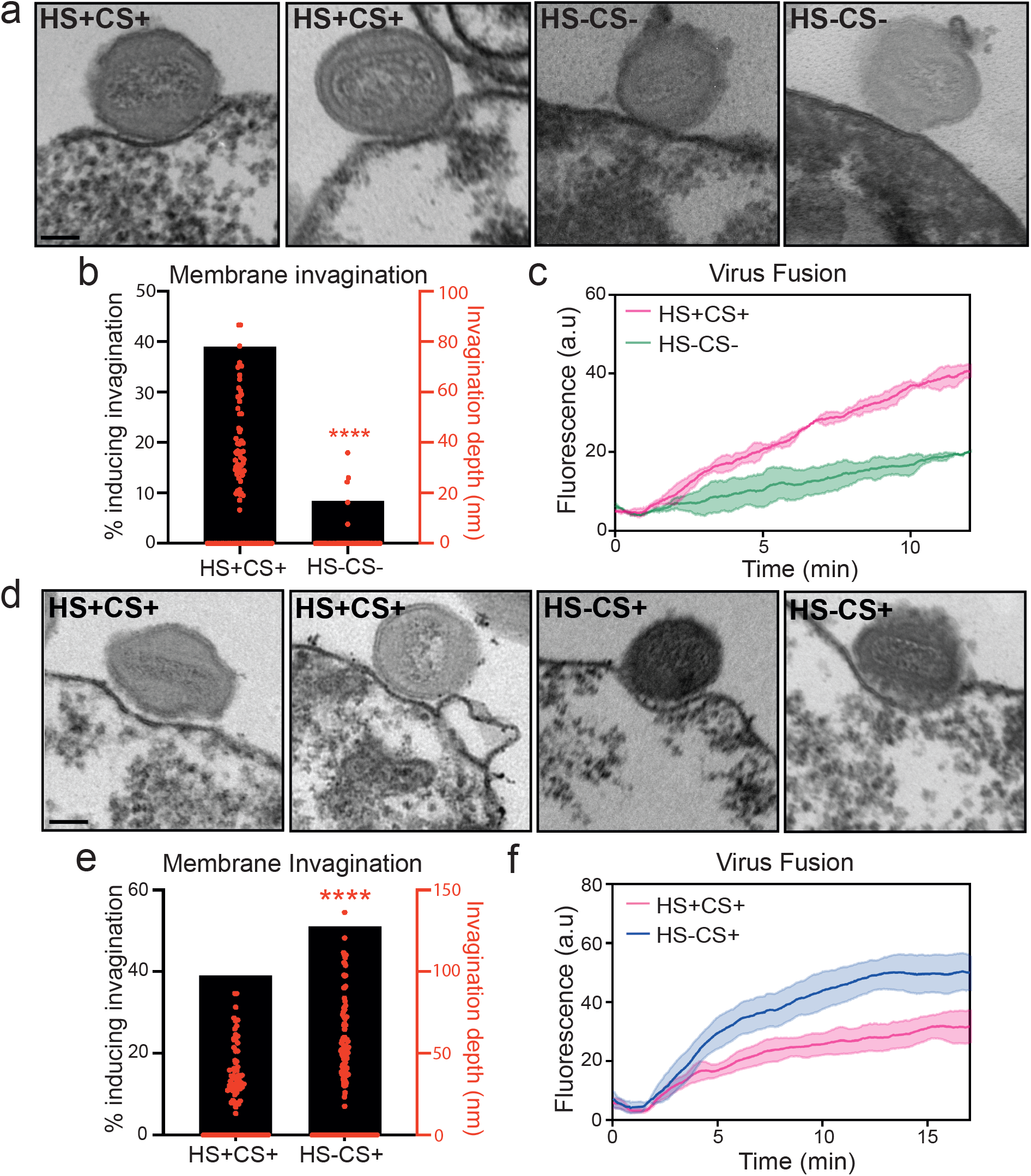
D8-mediated invagination requires chondroitin sulfate binding. **a.** TEM images of WT VACV bound to HS+CS+ or HS-CS- cell-derived blebs at pH 5.0. Scale bars = 100 nm **b.** Quantification of % invagination and invagination depth of virions from (**a**). **c.** Hemifusion rates of WT virus on HS+CS+ and HS-CS- cells using the R18 dequenching assay. **d.** TEM images of WT VACV bound to HS+CS + or HS-CS+ cell-derived blebs at pH 5.0. Scale bars = 100 nm **e.** Quantification of % invagination and invagination depth of virions in (**d**). **f.** Hemifusion rates of WT virus on HS+CS+ and HS-CS+ cells using the R18 dequenching assay. For **b** (n=70 virions/condition)**, c, e** (n=85 virions/condition) **and f** data are means ± SD. Statistical analysis was performed using unpaired twotailed *t*-tests (\*\*\*\**P* < 0.0001; ns, not significant (*P* > 0.05)).

Although H3 binds HS^11^, and our H3-binding experiments showed a significant reduction in VACV adsorption (Supplementary Fig. 2b), we saw only a minor reduction in VACV binding to cells lacking HS (Supplementary Fig. 4a). This was further corroborated with soluble GAG pre-incubation experiments. For this, A4-EGFP virions were pre-incubated with heparan, HS, CS-A, or CS-E, which was reported to be the specific binding partner of D8^36^, prior to adsorption onto HS+CS+ cells. Analysis of VACV binding by flow cytometry indicated that pre-incubation with heparan, HS or CS-A had little impact VACV attachment, while pre-incubation with CS-E blocked VACV binding by 30% (Supplementary Fig. 4b). When CS-E preincubation experiments were repeated on HS-CS+ cells we found that CS-E reduced binding to HS-CS+ cells by a further 27% relative to HS+CS+ cells (Supplementary Fig. 4c). These results indicate that CS-E is the predominant GAG used for VACV binding on HS+CS+ cells.

That VACV binding to HS-CS+ cells relied on CS-E to an even greater extent suggested to us that the D8/CS-E interaction may be compensating for the loss of H3-HS interactions and thereby masking any HS-binding phenotypes. To test this we compared the invagination depth of WT virions bound to HS+CS+ and HS-CS+ blebs at pH 5.0 using TEM. There was a notable increase in the depth of invagination under virions bound to HS-CS+ blebs compared to those bound to HS+CS+ blebs (Fig. 4d). Quantification showed that 12% more virions induced invagination on HS-CS+ cells and that the average depth of invagination was significantly increased (Fig. 4e). Consistent with a role for D8/CS-E mediated invagination in promoting hemifusion efficiency, R18-dequencing assays indicated that the rate of hemifusion was 2-fold higher on HS-CS+ cells than on HS+CS+ cells (Fig. 4f).

Together these results confirm CS-E as the cellular binding partner of D8 ^36^, define the importance of this interaction in assuring VACV fusion efficiency, and uncover the built-in compensatory nature of VACV cell surface binding proteins.

Having shown that VACV binding and fusion machineries are organized into distinct functional domains that dictate virion cell binding and fusion orientation^3^, we set out to characterize the relationship between side-on virion binding and tip-on virion fusion.

To do this we established a cell-derived bleb model system^24^. We show that blebs can be purified from multiple cell types and maintain the biologically relevant cell surface molecules needed to study virus binding and fusion.

Using blebs we determined that VACV side-on binding is static and that bound virions could undergo low-pH mediated hemifusion with blebs *i.e*. virions do not change orientation to bring the virus fusion machinery into contact with the target membrane. This led to the discovery that VACV virions can induce target membrane invagination in the presence of low pH. We found that invagination is mediated by the binding protein D8 and its interaction with the GAG CS. We show that this phenomenon is required for efficient low pH dependent membrane fusion activity at the plasma membrane and within macropinosomes; consistent with VACV entry by acid-mediated endocytosis ^4,5,30,39^.

Virus-induced membrane curvature has been described for SV40 in cells and GUVs and human norovirus on GUVs^19,21^. For both of these, interactions between pentameric virus scaffolds and glycosphingolipids are responsible for inducing plasma membrane curvature, and in the case of SV40 virus endocytosis^19^. The induction of membrane invagination seen here with VACV differs is several ways: it is mediated by virus-GAG interactions, is pH dependent, occurs in late endosomal compartments and it directly influences virus fusion efficiency.

Including this study, the low pH dependent processes that influence VACV entry are removal of the fusion suppressor A26^14^, protonation of the virus core^23^, D8-CS mediated invagination and virus fusion^4^. That removal of the fusion repressor and core activation can occur independent of target membrane contact suggests that these constitute the first low pH step, while D8-CS mediated invagination and virion fusion constitute the second.

Investigating hierarchy and redundancy between the VACV binding proteins using the four binding mutant VACVs in parallel, we found that A26>D8>H3 with regard to virion cell surface attachment and that A27 was not required. These findings are agreement with VACV ligand receptor capture experiments, which found VACV A26 (the laminin binding protein), a-dystroglycan (a laminin interacting protein) and CSPG4 (chondroitin sulfate proteoglycan 4) as significant interactions involved in VACV attachment to HeLa cells^40^. The degree to which VACV uses HS and CS for cellular attachment varies between experimental systems and cell types^15–18^. In partial explanation of this, our comparative studies on HS+CS+ and HS-CS+ cells indicate that VACV increased its usage of CS in the absence of HS. This suggests that VACV “adapts” attachment factor usage depending on which factors are available. This has also been seen with herpes simplex virus 1 (HSV-1), which adopts CS-E as a dominant attachment factor in the absence of HS^41^. In both cases, this attachment factor switching phenomenon likely acts to expand the cell type promiscuity and host range of these viruses.

We have previously suggested that “the organization of the VACV membrane into functionally distinct domains has evolved as a mechanism to maximize virion binding and fusion efficiency for productive infection”^3^. Here we demonstrate that VACV maximizes side-on binding and tip-directed virion fusion by directly linking these two processes through low pH dependent membrane invagination. In do so, we further demonstrate that virion protein architecture is critical to virus function and extend the role of virus binding proteins beyond that of mere attachment factors.

## Methods

### Cells Lines

BSC40, HeLa, L929, Gro2c and Sog9 cells were cultivated in Dulbecco’s modified Eagle’s medium (DMEM; Gibco, Life Technologies 11320033) supplemented with 10% heat-inactivated fetal bovine serum (FBS; Life Technologies 10500064), 2 mM Glutamax (Life Technologies 35050038) and 1% pen-strep (pen-strep; Sigma P0781).

### Viruses

Recombinant VACVs were generated in the Western Reserve (WR) strain. EGFP-A4^29^, A27(+/-) EGFP-A4^3^ and E-EGFP^42^ were previously described. ΔD8 and H3(-) were previously described as ΔD8vFire^4^ and vH3i^43^, respectively. ΔD8 EGFP-A4, H3(-) EGFP-A4 and WRΔA26 EGFP-A4 were constructed as previously described^23^. To generate ΔA26 EGFP-A4, A26 was deleted at its endogenous locus with mCherry, for ΔA26ΔD8 EGFP-A4, A26 was replaced with lacZ ΔD8 EGFP-A4. For HA-H3 or A26-HA, the N- and C-terminus were HA-tagged, respectively and recombinant virus selected for using E/L EGFP expression. Virus were selected and purified to homogeneity through four rounds of plaque purification and all resultant recombinant viruses confirmed by sequencing. All viruses were produced in BSC40 cells, and MVs band purified as previously described^5^. A27(-) and H3(-) virus stocks were generated in the absence of isopropyl β-d-thiogalactopyranoside (IPTG; Sigma 16758). Plaque forming units /ml and particle counts - calculated from the optical density at 260 nm^44^ - were determined for each purified recombinant VACV MV stock.

### Mature virion (MV) yield

Confluent 6-wells of BSC40, HS+CS+, HS-CS+ or HS-CS- were infected at a multiplicity of infection (MOI) of 10 for 1 hr, fed with full medium and infected cells collected at 24 hpi. Virus was released from cells by 3 rounds of freeze–thaw and plaque forming units/millilitre (pfu/ ml) determined by serial dilution of BSC40 cell monolayers.

### Antibodies

Anti-L1 mouse monoclonal antibodies (clone 7D11) was provided by B. Moss (National Institute of Health) with permission of A. Schmaljohn (University of Maryland). Anti-D8 rabbit polyclonal was provided by P. Traktman (Medical University of South Carolina). Anti-HA rabbit polyclonal antibodies were purchased from BioLegend (902302). Anti-A27 rabbit antibody (VMC39) was produced using purified recombinant baculovirus-expressed proteins by the Cohen lab^45^.

### Bleb Preparation

For bleb preparations, 3 confluent T75 flasks of cells were treated with 1.6 μM Latrunculin B (Sigma L5288) in DMEM for 15 min on an orbital shaker (700 rpm). The media was collected, detached cells pelleted (300 x g, 5 min) and the bleb-containing supernatant spun at 4,000 x g to pellet blebs. The bleb pellet was resuspended in buffer (IB; 10 mM NaCl, 280 mM k-glutamic acid, 14 mM Mg_2_SO_4_, 13.34 mM CaCl_2_, 5 mM Hepes) and filtered through a 5 μm filter (Cambridge Bioscience 43-10005-40). For ATP reconstitution, blebs were permeabilised with 50 mg/mL *Staphyloccus aureus* α-toxin (Hemolysin, Sigma H9395) and incubated with energy buffer (1 mM ATP, 1 mM UTP, 1 mM MgCl_2_, 10 mM creatine phosphate (Sigma 2380), 1 mg/mL creatine phosphokinase (Sigma 2384)) for 20 min at room temperature (RT), before pelleting and resuspending.

### Binding Orientation Assay

Blebs were labelled with CellMask Deep Red Plasma Membrane Stain (Invitrogen C10045) and pelleted (300 x g for 10 min) onto fibronectin (Sigma F1141) coated CELLview glass bottom cell culture slides (Greiner Bio-One 43079). A4-EGFP WT or mutant viruses were R18 labelled and bound to blebs at 4 °C for 1 hr. Samples were then washed and incubated with media adjusted with 100 mM 2-(*N*-morpholino)ethanesulfonic acid (MES) to pH 7.4 or pH 5.0 for 10 min at 37 °C. Samples were fixed with 4% formaldehyde 20 min, washed and imaged on an ELYRA PS.1 microscope (Zeiss).

### Structured Illumination Imaging

For viral membrane protein distribution, high-performance coverslips (18 × 18 mm, 1.5H, Zeiss) were washed and samples prepared and imaged as previously described ^27^.

### Polarity Factor

Polarity factors were calculated as in Gray *et al* ^3^.

### Flow Cytometry

Cells were seeded to confluency in 96-well plates (Greiner-Bio One 650101) and incubated with equal numbers of particles of each recombinant EGFP-A4 VACV strain for 1 hr at 4 °C. For cellular GAG and protein inhibition studies, virus was preincubated for 1 hr at 4 °C with 100 μg/mL soluble H (Sigma H4784), HS (Sigma H7640), CS-A (Sigma C9819), laminin (Sigma L6274) or CS-E (Cosmo Bio CSR-NACS-E2(SQC)3), pelleted, washed and the virus-added to cells for 1 hr at 4 °C. Unbound virus was removed by washing and cells fixed in 4% paraformaldehyde and harvested. Flow cytometry was performed on the Guava EasyCyte HT flow cytometer, recording the EGFP fluorescence with the 488 nm laser. Analysis of the flow cytometry data was performed with the GuavaSoft 3.3 analysis package (FlowJo).

### Electron Microscopy

For virus bound to blebs, blebs were centrifuged (700 x g for 10 min) onto fibronectin (Sigma F1141) coated CELLview glass bottom cell culture slides (Greiner Bio-One 543079). Virus was bound to blebs at 4 °C for 1 hr. Unbound virus was removed and samples incubated with media adjusted to either pH 7.4 or pH 5 with 100 mM MES for 10 min at 37 °C. Samples were fixed with 1.5% formaldehyde (Sigma F8775) 2% glutaraldehyde (Sigma 340855) for 20 min. For virus in macropinosomes, HeLa cells were grown to confluency on coverslips and recombinant virus bound at equal particle number at RT for 1 hr. Unbound virus was removed and full medium supplemented with 10 mg/mL horseradish peroxidase (HRP). Samples were shifted to 37 °C for 1 hr before fixation in 1.5% FA and 2% glutaraldehyde in 0.1 M cacodylate for 30 min. Samples were then incubated for 1 hr in 1% osmium tetraoxide/1.5% potassium ferricyanide at 4°C, treated with 1% tannic acid in 0.05 M sodium cacodylate for 45 min in the dark at RT, dehydrated in sequentially increasing concentrations of ethanol and embedded in Epon resin. The 70 nm thin sections were cut with a Diatome 45° diamond knife using an ultramicrotome (UC7, Leica). Sections were collected on 1×2 mm formvar-coated slot grids and stained with lead citrate. All samples were imaged using a transmission electron microscope (Tecnai T12, FEI) equipped with a charge-coupled device camera (SIS Morada, Olympus).

### TEM image analysis

For image analysis, invagination depth of bleb bound virions was quantified using the Olympus SIS iTEM software. A straight line was drawn across the two edges of the invagination and the perpendicular distance to the inner leaflet of the bleb membrane was determined, signifying the ‘invagination depth’. For macropinosome curvature analysis, the Kappa plugin^34^ in Fiji^46^ was used. In brief, an initialisation curve was traced using a point-click method along the macropinosome membrane in contact with the virion membrane. This was then fit to the underlying data using an iterative minimization algorithm. The Bezier curve was extracted and the mean curvature along the entire curve reported.

### Early gene expression (RT-qPCR)

RT-qPCR was performed as described in^47^. Briefly, total RNA was extracted from infected HeLa cells using the RNeasy Plus Mini kit (Qiagen 74134) according to manufacturer’s instructions. Amplification of the VACV early gene, C11 and (GAPDH) cDNA was performed by qPCR (Mesa Blue qPCR MasterMix Plus for SYBR assay; EurogentecB). Viral mRNA threshold cycle (CT) values are displayed as abundance normalized against GAPDH.

### Western Blot

Purified virions were heated at 95 °C for 10 min before separation on 4-12% Bis-Tris polyacrylamide gels (Thermo Fisher Scientific NW00125BOX) and transfer to nitrocellulose membranes. Membranes were blocked with 5% milk, 0.1% Tween-20 (TBST; Abcam ab64248) for 1 hr at RT. Membranes were incubated with primary antibody D8 or L1 (1:1000) and bands visualized with (HRP)- coupled secondary antibody (1:5,000) (Cell Signalling, rabbit 7074S, mouse 7076S) on an ImageQuant LAS 4000 Mini (GE Life Sciences) with Luminata Forte Western HRP Substrate (Sigma WBLUF0500) for detection.

### Bulk Fusion Measurements

Bulk fusion was performed as described in Schmidt *et al*^23^. Breifly, Equal MV particles were labelled with R18 (R18, ThermoFisher Scientific O246) and bound to HeLa cells or blebs and the pH maintained at 7.4 or lowered to 5.0 by the addition of 100 mM MES.. After acquisition, total R18 was dequenched by the addition of 10% Triton X-100. R18 fluorescence was normalised to the signal intensity after the addition of Triton X-100. R18 fluorescence was measured using a Horiba FluoroMax-4 (Horiba Jobin Yvon) spectrofluorometer with an excitation wavelength of 560 ± 5 nm and an emission wavelength of 590 ± 5 nm.

### Data Availability

The datasets generated and/or analysed during the current study are available from the corresponding author on reasonable request.

## Acknowledgements

We thank B. Moss for providing the mutant viruses that were used in this study. Laura Pokorny is supported by the UCL Birkbeck MRC DTP. J.J.B. is supported by MRC core funding to the MRC Laboratory for Molecular Cell Biology at University College London, award code (MC_U12266B). D.A. was funded by a Marie Skłodowska-Curie fellowship funded by the European Union (750673), R.B. was funded by the MRC Laboratory for Molecular Cell Biology PhD program. K. E.L. and Y.M are supported by a Wellcome Trust Senior Research Fellowship (217191/Z/19/Z) to Y.M., P.S. and T.J.K. are supported by BBSRC grants (BB/P0098401 & BB/S017283/1) to T.J.K., and J.M. was supported by the European Research Council (649101-UbiProPox) to J.M and core funding to MRC Laboratory for Molecular Cell Biology (MC_UU_00012/7) to J.M.

## Author contributions

L. P. and J.M conceived the project. All authors contributed experimental design. L.P, J.J.B., D.A., R.B., K.E.L., and P.S. performed the experiments. L.P. and J.M. and wrote the manuscript. L.P. and J.J.B performed the EM. All authors performed aspects of data analysis. L.P, Y.M., T.J.K and J.M discussed the results and implications of the findings. All authors discussed the manuscripts provided comments and editing.

## Competing Interests

The authors declare no competing interests.

## Supplementary Information

**Supplementary Figure 1.**
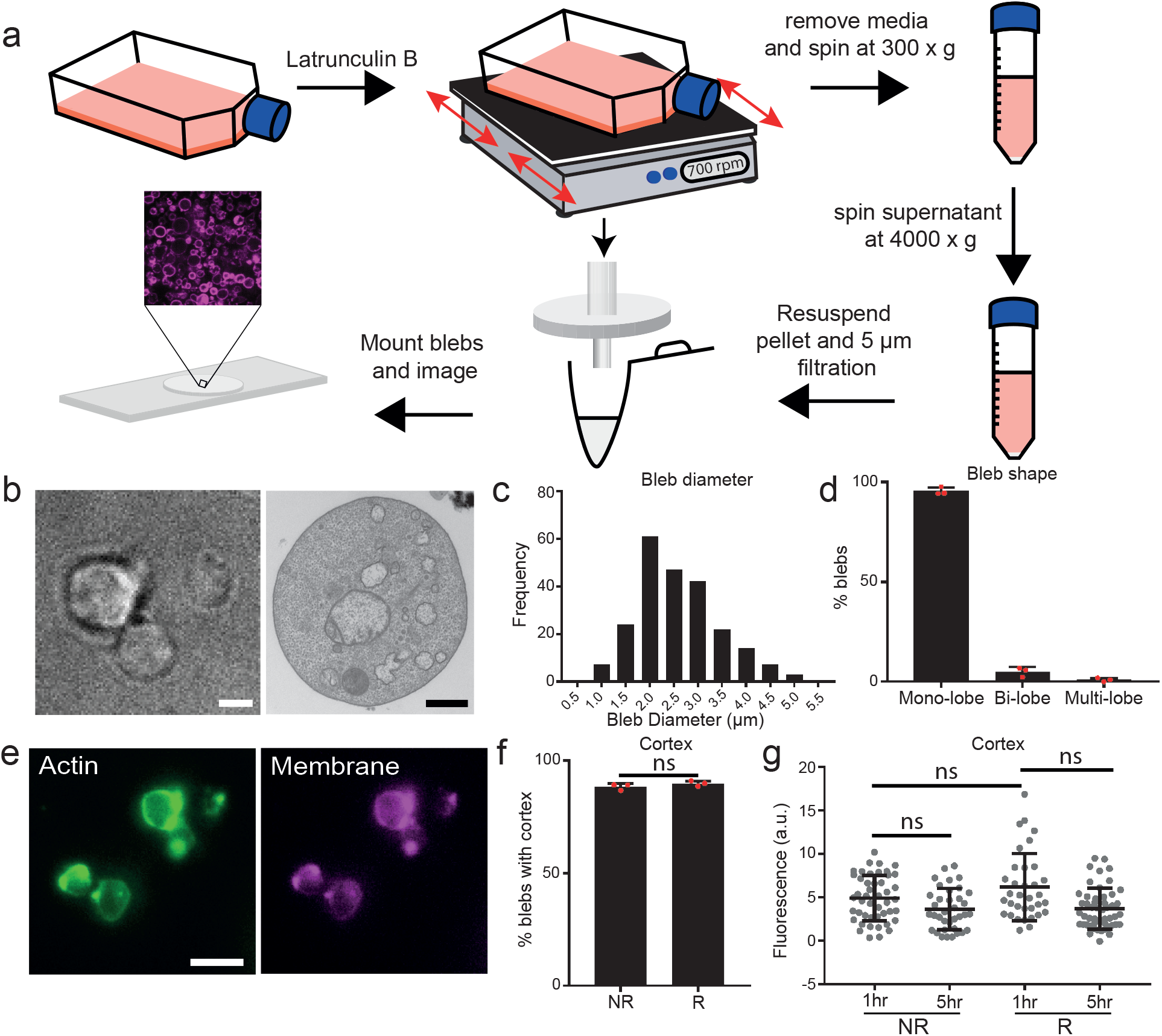
Cell-derived membrane blebs as a minimal cell system. **a.** To prepare blebs Latrunculin B is added to cells to induce blebbing. Cells are shaken to detach blebs, cellular debris removed by a slow spin step (300 x g), blebs collected by a fast spin step (4,000 x g), and remaining large debris removed by filtration through a 5 μm pore filter. **b.** Representative brightfield (LHS, scale bar; 1 μm) and TEM (RHS, scale bar; 500 nm) images of blebs after purification. **c.** Histogram of bleb diameter range. **d.** Blebs were scored for mono-, bi- and multi-lobulation. **e.** Blebs were stained for actin (green) and the plasma membrane (PM; magenta). Scale bar, 5 μm. **f.** Bleb cortical actin was reconstituted (R) with ATP or not (NR), stained for both actin and the PM and the percentage of blebs with an actin cortex calculated. **g.** Stability of the actin cortex over time at 37 °c between NR and R blebs was determined by intensity measurements of the actin stain on z-projections. For **d, f** and **g** data are means ± SD. Statistical analysis was performed using unpaired two-tailed *t*-tests (ns, not significant (*P* > 0.05)).

**Supplementary Figure 2.**
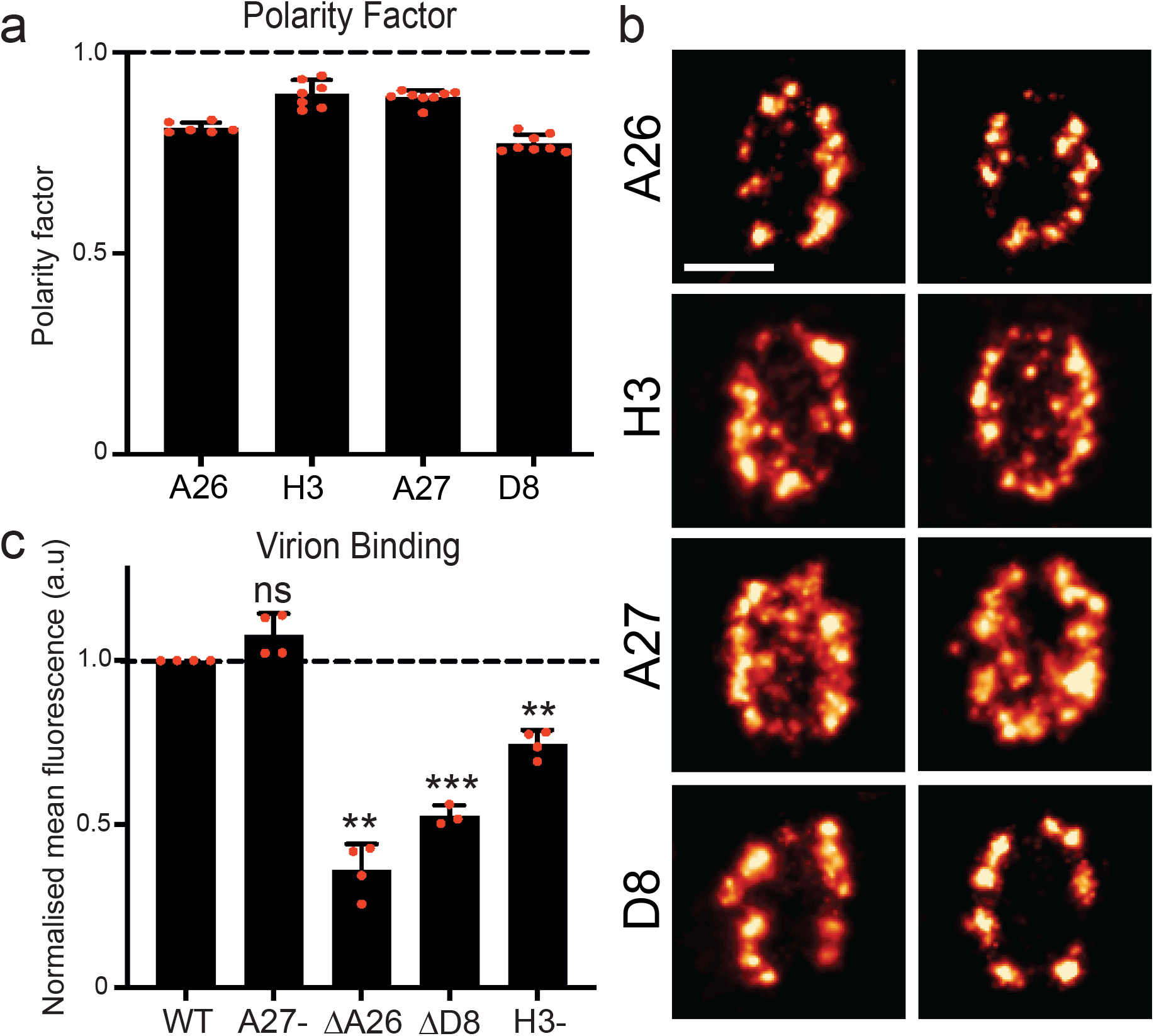
VACV binding proteins reside at virion sides and are differentially required for VACV binding. **a.** Quantification of VACV binding protein polarity factors using data from the models in Fig. 2A. A polarity factor of less than one corresponds to concentration of the protein at the sides of MVs. **b.** Representative STORM images of VACV binding proteins on individual MVs. Scale bar = 200 nm. **c.** Binding affinities of WT and recombinant binding protein mutant VACVs on HS+CS+ cells. For **a** and **c** data are means ± SD. Statistical analysis was performed using unpaired two-tailed *t*-tests (\*\*\**P* < 0.001; \*\**P* < 0.01; ns, not significant (*P* > 0.05)).

**Supplementary Figure 3.**
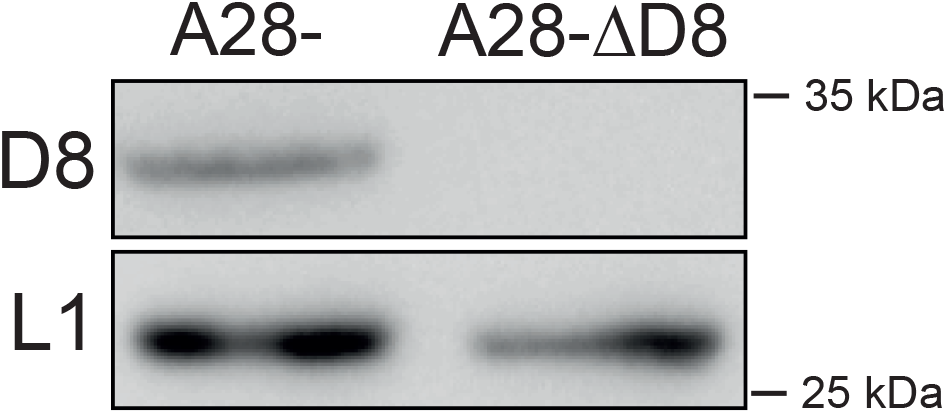
Confirmation that A28-ΔD8 VACV virions do not package D8. Representative immunoblot of D8 protein packaging in A28- and A28-ΔD8 virions. Molecular weight markers are indicated at right.

**Supplementary Figure 4.**
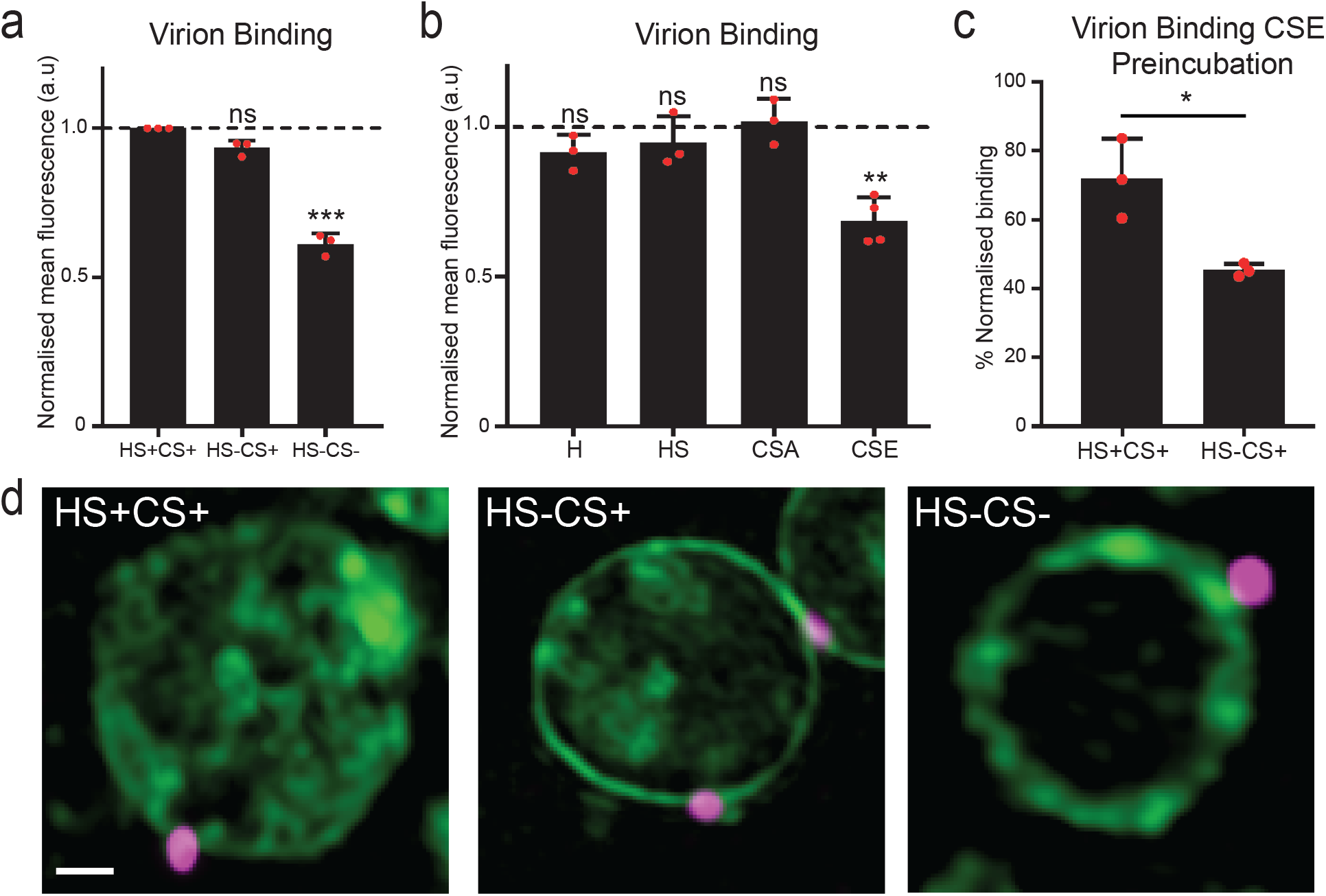
CS-E is the major GAG used by VACV for binding. **a.** Binding affinities of WT VACV on HS+CS+, HS-CS+ and HS-CS- cells. **b.** Binding affinities of WT VACV with GAG preincubation on HS+CS+ cells. **c.** Binding affinities of WT virus with CS-E preincubation on HS+CS+ and HS-CS+ cells. Data is normalised to no CS-E preincubation on the given cell type. **d.** SIM images of VACV bound to HS+CS+, HS-CS+ and HS-CS- derived blebs scale bar = 1μm. A4-EGFP VACV (magenta) and PM (green). For **a, b** and **c** data are means ± SD. Statistical analysis was performed using unpaired two-tailed *t*-tests (\*\*\**P* < 0.001; \*\**P* < 0.01; \**P* < 0.05; ns, not significant (*P* > 0.05)).

## Notes

### Competing Interest Statement

The authors have declared no competing interest.

